# Informatics investigations into anti-thyroid drug induced agranulocytosis associated with multiple HLA-B alleles

**DOI:** 10.1101/713743

**Authors:** Kerry A Ramsbottom, Daniel F Carr, Daniel J Rigden, Andrew R Jones

## Abstract

Adverse drug reactions have been linked with HLA alleles in different studies. These HLA proteins play an essential role in the adaptive immune response for the presentation of self and non-self peptides. Anti-thyroid drugs methimazole and propylthiouracil have been associated with drug induced agranulocytosis (severe lower white blood cell count) in patients with B*27:05, B*38:02 and DRB1*08:03 alleles in different populations: Taiwanese, Vietnamese, Han Chinese and Caucasian.

In this study, informatics methods were used to investigate if any sequence or structural similarities exist between the two associated HLA-B alleles, compared with a set of “control” alleles assumed not be associated, which could help explain the molecular basis of the adverse drug reaction. We demonstrated using MHC Motif Viewer and MHCcluster that the two alleles do not have a propensity to bind similar peptides, and thus at a gross level the structure of the antigen presentation region of the two alleles are not similar. We also performed multiple sequence alignment to identify polymorphisms shared by the risk but not by the control alleles and molecular docking to compare the predicted binding positions of the drug-allele combinations.

Two residues, Cys67 and Thr80, were identified from the multiple sequence alignments to be unique to these risk alleles alone. The molecular docking showed the poses of the risk alleles to favour the F-pocket of the peptide binding groove, close to the Thr80 residue, with the control alleles generally favouring a different pocket. The data are thus suggestive that Thr80 may be a critical residue in HLA-mediated anti-thyroid drug induced agranulocytosis, and thus can guide future research and risk assessment.

## Introduction

Adverse drug reactions have been linked to Human Leukocyte Antigens (HLA) in multiple different studies, where an individual carrying a specific risk allele has a higher risk of developing a reaction to that drug, including skin conditions like Stevens-Johnson syndrome and toxic epidermal necrolysis (SJS/TEN) and drug induced liver injury (1–3). HLA proteins play a role in the adaptive immune response, presenting peptides to the T-cell receptors. Non-self peptides are then recognised and elicit an immune response where appropriate (4). Occasionally, unnatural interaction of drugs during this process results in an adverse drug reaction. The role of HLA in these adverse drug reactions has been hypothesised in three main ways: the Hapten model, the Pharmacological Interaction model and the Altered Peptide Repertoire model. The *Hapten* model predicts the drug binds covalently to a self-protein and is processed via HLA molecules; this drug-protein combination is presented and recognised as being non-self, initiating an immune response (5). The *Pharmacological Interaction (PI)* model predicts the drugs bind directly to the TCR or via the formation of HLA-drug complexes which activate T cells and thus initiate an immune response without the need for a specific peptide ligand (6). Except where noted below, the results presented here work under the assumption that the drugs we are investigating here will follow the *Altered Peptide Repertoire* model, with the drug interacting non-covalently with the HLA molecule directly within the antigen presentation site. This leads to a difference in the self-peptide set that is presented to the T-cell receptors and thus, initiating an immune response (7). Currently, the most widely investigated HLA-ADR association is that of abacavir with B*57:01. For this association, the crystal structure of the drug bound in complex with the risk allele is available (8, 9). Illing *et al.* (8) and Ostrov *et al*. (9) have demonstrated the Altered Peptide Repertoire model with high confidence for abacavir, including the crystal structure for abacavir bound in the peptide binding groove of the associated risk allele, B*57:01, along with proteomics evidence for different peptides being presented in the bound and unbound cases (8).

The peptide binding groove of the HLA is a long hydrophobic cleft formed between the α-helices and β-sheet platform. This cleft is much larger than the naturally involved binding sites that proteins have for small organic molecules. The peptide binging groove contains six subsites (S4 Fig). The size and stereochemistry of the subsites are determined by the polymorphic residues along the cleft (10). The specificity of peptide binding is determined, in part, by the interactions between anchor residues on the peptide side chains at two or more of these subsites (11).

Anti-thyroid drugs are used to treat hyperthyroidism as they normalise thyroid function through binding to the thyroid peroxidase enzyme (12). These drugs are thioamides containing a thiocarbonyl group and a thiourea moiety within a heterocyclic structure. The common agents used are methimazole, carbimazole and propylthiouracil (13). Agranulocytosis has been defined as absolute neutrophil count below 500/µl of blood and includes not only neutrophil count but also the absolute number of eosinophils, basophils and mast cells (14). Patients with severe neutropenia are likely to experience infections which may be life-threatening or even fatal. The mechanism of anti-thyroid induced agranulocytosis is through either direct toxicity or immune-mediated toxicity (14). The incidence of agranulocytosis in England and Wales has been estimated at around 7 cases per million people per year, with an adjusted odds ratio for neutropenia of 34.7 for users of thyroid inhibitors (i.e. anti-thyroid drugs) (15). Most cases of agranulocytosis are idiosyncratic reactions to drugs or their metabolites, including anti-thyroid drugs. Other causes include splenic sequestration, nutritional deficiencies, infections, immune neutropenia, haematological disease and primary congenital or chronic neutropenia (16). An increased risk of agranulocytosis has been associated with anti-thyroid drugs carbimazole, methimazole and propylthiouracil for three different HLA alleles in Asian and Caucasian populations, these associations are shown in Table 1.

**Table 1:**
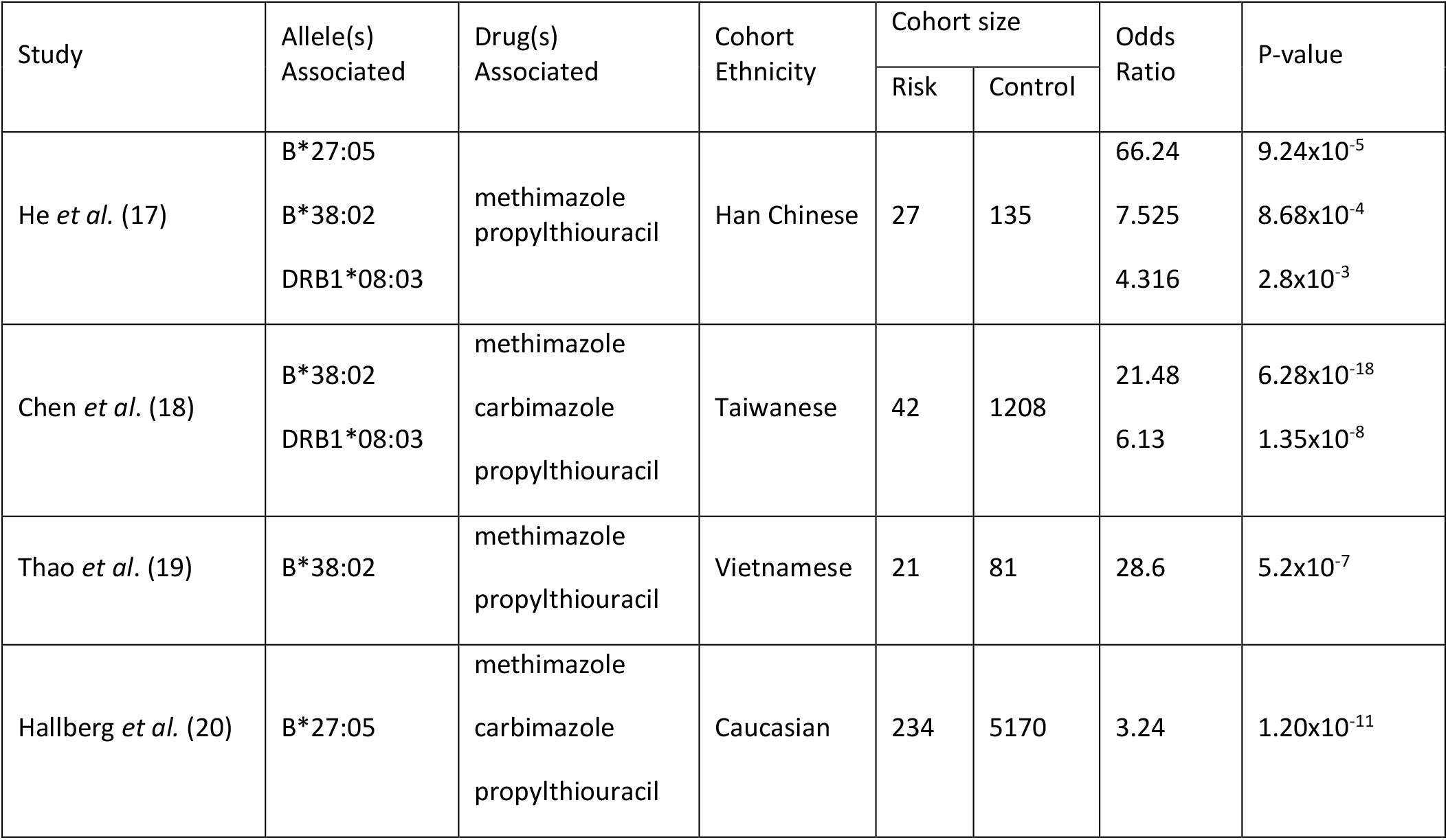
HLA associations seen carbimazole, methimazole and propylthiouracil across populations.

| Study | Allele(s)<br>Associated | Drug(s)<br>Associated | Cohort<br>Ethnicity | Cohort size |  | Odds<br>Ratio | P-value |
| --- | --- | --- | --- | --- | --- | --- | --- |
|  |  |  |  | Risk | Control |  |  |
| He *et al.* (17) | B*27:05 | methimazole<br>propylthiouracil | Han Chinese | 27 | 135 | 66.24 | $9.24 \times 10^{-5}$ |
| | B*38:02 | | | | | 7.525 | $8.68 \times 10^{-4}$ |
| | DRB1*08:03 | | | | | 4.316 | $2.8 \times 10^{-3}$ |
| Chen *et al.* (18) | B*38:02 | methimazole<br>carbimazole<br>propylthiouracil | Taiwanese | 42 | 1208 | 21.48 | $6.28 \times 10^{-18}$ |
| | DRB1*08:03 | | | | | 6.13 | $1.35 \times 10^{-8}$ |
| Thao *et al.* (19) | B*38:02 | methimazole<br>propylthiouracil | Vietnamese | 21 | 81 | 28.6 | $5.2 \times 10^{-7}$ |
| Hallberg *et al.* (20) | B*27:05 | methimazole<br>carbimazole<br>propylthiouracil | Caucasian | 234 | 5170 | 3.24 | $1.20 \times 10^{-11}$ |

The structures of methimazole and propylthiouracil are shown in S5 Fig. It can be seen that these two associated drugs share a common thiocarbonyl group, which is also seen in the experimental anti-thyroid drugs. The associated anti-thyroid drugs share a common target, thyroid peroxidase, in their normal mechanism of action (21). A study by Pradhan *et al.* conducted molecular docking of methimazole and propylthiouracil with thyroid peroxidase. The results of this study predicted both drugs to bind in the same position, with the sulphur group forming a hydrogen bond with Arg491 of the thyroid peroxidase, resulting in inhibition of thyroid hormone production (12). As these drugs show similar binding to the same target, it is reasonable to hypothesise that both drugs might also have a shared mechanism for the adverse drug reaction, going some way to explaining why multiple drugs have been associated with the same alleles. It can be noted that methimazole is more commonly associated than propylthiouracil, although neither drug has been studied in isolation (17, 18, 20). In the He et al. study, there were 26 methimazole agranulocytosis cases compared to 3 for propylthiouracil. For the Chen et al. study, of the 42 thioamide-induced agranulocytosis cases, it was stated that 9 of these were patients taking carbimazole, 9 propylthiouracil and 23 methimazole. For the Hallberg et al. study, 39 cases were induced by anti-thyroid agents, of these 29 were methimazole, 5 carbimazole and 5 propylthiouracil. Although all three anti-thyroid drugs have been incorporated into the association studies, our study focuses on the associations seen with methimazole and propylthiouracil. Carbimazole is the pro-drug of methimazole, responsible for the antithyroid activity, and has a short half-life of 5.3-5.4 hours with peak plasma concentrations of methimazole being present after 1 or 2 hours (22, 23). We are working under the assumption that the mechanisms involved in the adverse drug reaction follow the altered peptide repertoire model, the active drug methimazole would be most likely to interact with the HLA during this process and so carbimazole is excluded from this analysis.

B*38:02 has a very similar sequence to B*38:01 with only one residue difference (B*38:02 Thr80Ile B*38:01). It therefore is highly possible that B*38:01 could also be a risk allele in this context. B*38:02 (S6 Fig) is most commonly found in Asian populations (0.69%-6.6%, frequencies taken from Allele Frequency Net Database (AFND) (24)), with Caucasian (0.003-0.2%) and African populations (0.007-0.06%) having much lower frequencies. B*38:01 is more commonly found in Caucasian populations (0.6-6.7%), compared to Asian populations (0.15-1.0%) and therefore allele frequencies for B*38:01 in the Asian populations studies are likely not detected (S7 Fig). The Hallberg *et al*. study (which found B*27:05 to be significantly associated) was based on Caucasian populations with a high proportion of Swedish patients. Comparing the allele frequencies from AFND (24) for B*27:05 (S8 Fig) and B*38:01 in Swedish populations, B*27:05 shows frequencies between 10.5 and 20% with B*38:01 showing no entries – usually indicating that the allele was not detected in these populations. It could therefore be plausible that the B*38:01 allele could be an ADR-associated allele but this is not seen in the association study due to B*38:01 being a low frequency allele in Swedish populations. However, without case-tolerant association studies showing patients with B*38:01, this association cannot be confirmed or rejected. For the purpose of this study, the B*38:01 is considered as a possible risk allele.

A previous study conducted by Chen *et al.* completed molecular docking with methimazole and propylthiouracil docked with B*38:02, B*38:01 and DRB1*08:03. This study showed poses favouring both the B- and F-pockets along the peptide binding groove. Four suspected key residues were identified: Cys67, Asn77, Thr80 and Thr123. Their study focussed on the associations seen in Taiwan individuals and therefore does not include investigations into the B*27:05 alleles seen in Caucasian populations. Due to the differences in general structure between Class I and Class II HLA, it is difficult to compare docking predictions between the two allele classes. The B*38:02 and B*38:01 alleles were modelled using the same template structures, with the DRB1*08:03 structure modelled using only DRB1*01:01 as a template. The quality of the models used for molecular docking can impact the docking results and therefore the selection of templates is very important.

The purpose of this study is to investigate the associated alleles, in particular looking for commonalities between the two associated HLA-B alleles in order to look for similarities in these alleles that might shed light on the underlying mechanisms of the adverse drug reactions seen.

## Methods

### Sequence and structural analysis

In order to conduct a reliable analysis between cases and controls, the control alleles must first be carefully selected to ensure they can be reliably assumed to be non-associated. The case and control frequencies from He *et al.* (17), Chen *et al.* (18) and Hallberg *et al.* (20) were compared alongside healthy individual frequency data obtained from AFND (24). Alleles where the study control allele frequency or healthy individual frequencies sourced from AFND were similar to or greater than the case allele frequencies were selected as controls (Supplementary material i). These alleles could safely be assumed to be non-associated as they do not show enrichment in the case groups.

Structural differences were assessed between the risk and selected control alleles. Firstly, the peptide binding regions were compared, this is likely where the drug binding would occur and so differences here would be important for the mechanism of interaction involved in the adverse drug reaction. The peptide binding regions were compared using MHC motif viewer (25) and MHCcluster (26). MHC motif viewer was used to compare the predicted binding motifs for the B*27:05 and B*38:02 risk alleles as well as the B*38:01 possible risk and the selected control alleles. MHCcluster was similarly used to compare the global similarities of peptide binding predictions for the risk, possible risk and selected control alleles. Both the MHC motif viewer and MHCcluster post-process NetMHCpan scores to give predictions of motifs or similarities between peptides. NetMHCpan uses artificial neural networks to predict the peptide binding of HLA molecules based on IEDB experimental data of peptides known to be presented by given alleles, including data for the binding of peptides to B*27:05 and B*38:01 (27, 28). Therefore, the predictions generated will be based largely on experimental data.

The protein molecules were further compared using multiple sequence alignment to view differences across the whole protein and also individual residue changes within sub-pockets along the peptide binding groove between the risk and control alleles. Firstly, the sequences for the risk, possible risk and control alleles were aligned. The alignments were then extended to look at common alleles selected from AFND and NMDP (National Marrow Donor Program) (20, 24) Asian and Caucasian populations and extended further again to consider common alleles found in all populations. When considering the common alleles, it is important to note that the definitive risk profile (association status) of these alleles is unknown. Although, due to the high frequency and the fact that they have not been seen to be associated with agranulocytosis, it can be assumed that these alleles are likely not associated. These multiple sequence alignments allow us to look for residue similarities unique to the risk alleles and therefore identify residues which may be involved in the mechanism of binding for the adverse drug reactions.

### Molecular Docking

Molecular docking was used to compare the predicted binding sites between risk and control alleles. Crystal structures were obtained from the PDB database where available. For those alleles where the crystal structures are not available, the structures were predicted using Modeller (29, 30). Crystal structures, given the suffix ‘_S’, were available for B*27:05 (1OGT (31)), B*15:01 (1XR9 (32)) and B*51:01 (1E27 (33)). Homology models, given the suffix ‘_M’, were created for B*38:02, B*38:01, B*40:06, B*46:01 and B*54:01 (S1 Table). Methimazole, the active form of carbimazole, and propylthiouracil were used to dock with the B*27:05_S and B*38:02_M risk alleles, B*38:01_M possible risk allele and selected control alleles; B*15:01_S, B*40:06_M, B*46:01_M, B*51:01_S and B*54:01_M.

Structures and sequences of three similar alleles were obtained searching the PDB database using BLAST-P (34). The structures for these similar alleles, with high sequence identity (e.g. 95-98% identity for the B*38:02 templates), were then used as templates for Modeller. Target and template sequences were aligned with ClustalX (35) and ten models were made for each structure, using Modeller 9.9 automodel class (36). The model with the lowest objective function was chosen for the docking. S1 Table summarises the structures and models obtained. Drug structures for methimazole and propylthiouracil were both obtained from the PDB database (5FF1 (37) and 5HPW (38) respectively).

AutoDockFR (39) was used to dock both methimazole and propylthiouracil with the B*27:05 and B*38:02 risk alleles, B*38:01 possible risk allele and the selected control alleles. Structural PDBQT files were prepared using AutoDock Tools (40) for both the alleles and drug structures. AutoGrid (41) was used to map the target allele structures and select grid points in order to search both the peptide binding groove (PBG) and the top three largest pockets (Top3). The top three largest pockets were selected by pocket volume. Searching the largest pockets on the protein increases the search space to cover more of the protein and allows an alternate binding position away from the peptide binding region. This helps identify if the peptide binding groove is in fact the most favourable binding region, ensuring the most favourable poses are obtained. A total of 10 poses were obtained for each allele for each drug and each search space, resulting in 20 poses per allele for each drug. Further analysis was completed conducting 100 runs for each drug-allele combination (supplementary material ii), this allowed further analysis of the favouring of binding pockets. These poses were then automatically assigned positions (i.e. binding in B or F pocket) using k-mean analysis and used to visualise the favouring of binding positions for each drug-allele combination. LigPlot (42) was used to visualise the interactions for each of the lowest scoring poses for each drug-allele combination, in order to compare the residues involved with binding.

In order to investigate how the size and structure of the ligands and pockets could be having an impact on the molecular docking results, similar molecules were docked to each of the risk and control alleles. The PDB database was searched for ligands with ≥50% similarity to methimazole and propylthiouracil. Four ligands were selected: MZY (1,3-dihydroimidazole-2-thione) and TUL (2-thioxo-2,3-dihydropyrimidin-4(1H)-one) have been used as experimental antithyroid drugs and include a thiocarbonyl group, DMI (2,3-Dimethylimidazolium Ion) and EV0 (2-amino-6-propylpyrimidin-4(3H)-one) do not include the thiocarbonyl group, (S5 Fig). These structures were prepared in the same way as the associated anti-thyroid drugs, using AutoDock Tools (40). AutoDockFR (39) was used to dock the non-associated compounds with B*27:05 and B*38:02 risk alleles, B*38:01 possible risk allele and selected control alleles. The binding positions of these were then compared to those of the associated drugs, in order to deduce if the size and structures of the ligands and pockets could be having an impact on the molecular docking results.

## Results

### Sequence and Structural Analysis

We firstly investigated whether allele frequencies for controls were representative of larger populations available in similar regions. This allows us to test for potential biased sampling in source studies, especially as controls have been combined from different countries, as well as to determine our own control (non-associated) alleles for further comparison. Case and control frequencies were calculated from the data provided for the He *et al.* and Hallberg *et al*. studies (17, 20). Alleles with study control frequency over 3% were investigated. Five alleles were selected to be used as controls: B*15:01, B*40:06, B*46:01, B*51:01 and B*54:01. It can be reasonably assumed that these alleles are non-associated alleles based on the frequency data of both the Han Northern China and European populations (S1 Fig). The control allele selection is covered in more detail within the supplementary material (i).

The MHC Motif viewer (25) displays the preference for given HLA alleles to bind peptides with amino acids at given positions within an n-mer peptide e.g. 9mer for HLA class I. The motifs have been generated via the NetMHCpan (27) prediction method being run over a large selection of natural peptides, which has been trained originally with experimental data (peptides presented by given HLA alleles) from the IEDB database (28). While the predicted motifs cannot give us a direct measure of the likelihood of a drug to bind in the cleft of a given HLA allele, they can be indicative of whether different alleles share similar peptide binding regions. If the hapten hypothesis of HLA-mediated ADRs was true for anti-thyroid drugs, then the peptide binding ability of associated alleles might be related, with the caveat that drugs binding to peptides would likely change their affinity for particular alleles. Comparing the peptide binding motifs from MHC Motif Viewer (S9 Fig) it can be seen that although, as expected due to the high sequence similarity, the B*38:02 and B*38:01 alleles show very similar binding motifs, the B*27:05 allele shows a very different motif. This shows that there are differences between the favoured peptides and thus the peptide binding grooves of the B*27:05 and B*38:02 risk alleles. A similar observation can be made from the output for MHCcluster (Fig 1), with differences seen between B*27:05 and B*38:02. MHCcluster is also based on predictions from random human peptides processed by NetMHCpan and generates a distance measure between the peptide binding specificity scores generated by the software to create a tree representation. Similar to MHC Motif Viewer, the results can tell us about overall relatedness of predicted peptide binding by different alleles, but not directly about the likelihood of drug binding. From the tree-based output, it can be seen that the B*38:02 and B*38:01 alleles show very similar clustering, whereas the B*27:05 allele shows clustering differing from this and is not more closely related to the other risk allele than any of the selected control alleles. The differences seen between the favoured peptides of the risk alleles allows us to conclude that these alleles are not structurally similar.

**Fig 1:**
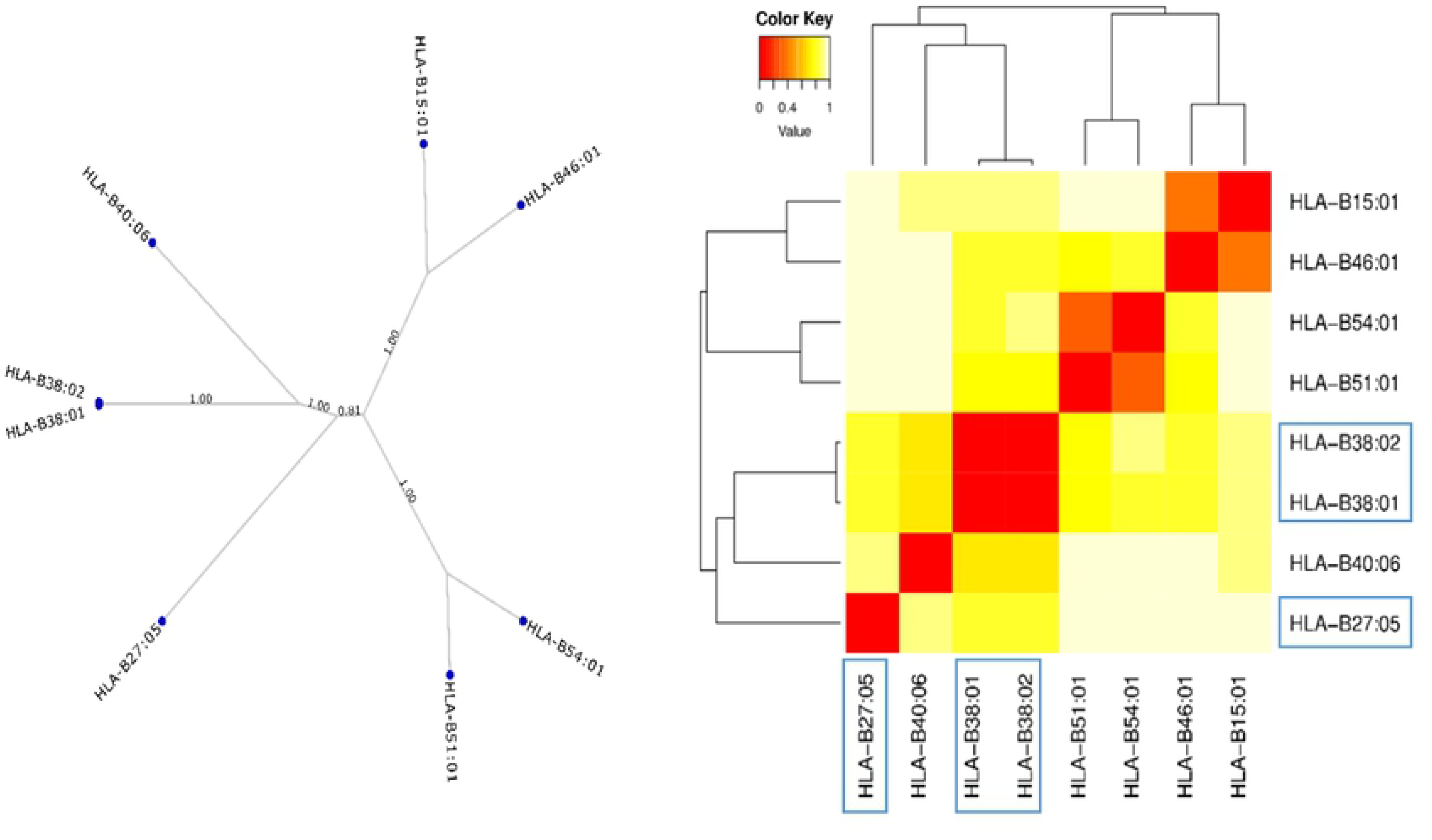
MHCcluster output for risk and control alleles. Specificity tree shows clustering of alleles and heat map shows the similarity between the binding motifs of each B allele, comparing known risk and possible risk (B*38:01) alleles (highlighted in blue) with selected controls. The scale shows the distance between the alleles, with red (0) showing very similar binding motifs and white (1) showing dissimilar binding motifs. Trees to the left and above the matrix show the hierarchical clustering of the different B alleles.

Multiple sequence alignments were used to investigate individual residue differences between alleles. Comparing the B*27:05 and B*38:02 risk alleles with the B*38:01 possible risk and selected controls, it can be seen that there are two residues which are seen to be unique to the risk alleles: Cys67 and Thr80 (Fig 2). These are both residues that were identified as potentially important in the Chen *et al.* study (18). When the comparisons were extended to look at common alleles, obtained from searching AFND for Caucasian and Asian populations and also common alleles found in the NMDP database (20) (S10 Fig), a similar pattern can be seen. These two residues are rarely found in these common alleles, although it must be noted that the association status of these common alleles is unknown.

**Fig 2:**
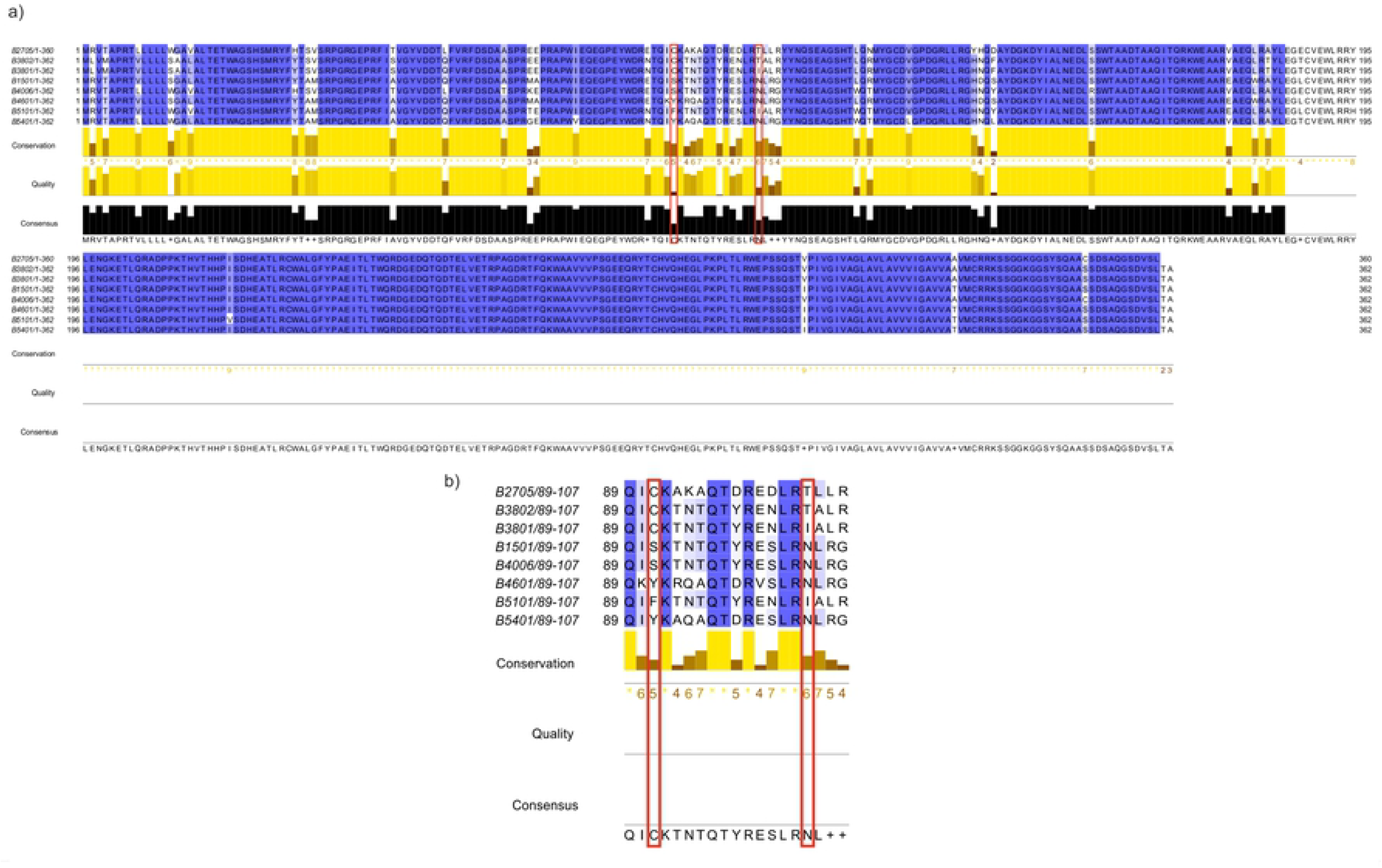
Multiple sequence alignment for risk and control alleles. (a) Multiple sequence alignment for risk alleles B*27:05 and B*38:02, possible risk allele B*38:01 and selected controls B*15:01, B*40:06 B*46:01, B*51:01 and B*54:01. (b) Focusing on positions 65-83. Positions 67 and 80 highlighted in red.

## Molecular docking

### Methimazole

Methimazole was docked with each of the risk, possible risk and selected control alleles, using AutoDockFR (39), in order to compare the favourable binding positions in each case. Table 2 summarises the poses seen for methimazole, showing the pocket of the lowest scoring pose, the number of poses in each pocket and the median scores for the poses in those pockets, for each search space. From this and Fig 3, showing the predicted binding positions for the top scoring pose for each allele, it can be seen that the risk alleles favour the F-pocket for drug binding, close to the position 80 identified. Both scores as well as the number of poses in each pocket, while searching the peptide binding region, favour this pocket. Extending the search space to cover the top three largest pockets, B*27:05_S favours a pocket outside of the peptide binding groove, although still close to the position 80 identified. B*38:02_M still favours the F-pocket and B*38:01_M favours the B-pocket, with some poses seen outside of the peptide binding groove. Control alleles B*15:01_S, B*46:01_M and B*51:01_S all favour the B-pocket, both with scores and number of poses, while B*40:06_M and B*54:01_M favour the F-pocket, searching the peptide binding groove. Extending the search space to cover the top three largest poses for each of these alleles, B*15:01_S now favours a pocket outside of the binding groove, B*46:01_M still shows favouring of the B-pocket and B*40:06_M, B*51:01_S and B*54:01_M favour the F-pocket. Comparing the interactions seen from the LigPlot diagrams (S11 Fig), it can be seen that B*27:05_S and B*54:01_M both are predicted to have potential interactions with methimazole at residue 80. The residue at position 80 of B*27:05_S forms hydrophobic interactions with the thiocarbonyl group of the methimazole, where the position 80 residue for B*52:01_M interacts with one of the nitrogen atoms.

**Table 2:** Summary of molecular docking positions for methimazole. Position of the lowest scoring methimazole pose along with the number of poses in each pocket and the median of the pose scores in each pocket for each of the alleles using the search space covering the peptide binding groove and the top 3 pockets identified on the protein (Extended search space). Scores given as kcal/mol. ‘O’ refers to pockets other than the B and F, i.e. outside of the binding groove.

| Status | Allele | Peptide Binding Groove |  |  |  |  | Extended Search Space |  |  |  |  |  |  |
| --- | --- | --- | --- | --- | --- | --- | --- | --- | --- | --- | --- | --- | --- |
|  |  | Lowest | B | Median | F | Median | Lowest | B | Median | F | Median | O | Median |
| Risk | B*27:05_S | F | 5 | -3.20 | 5 | -3.26 | O | 2 | -3.15 | 0 | N/A | 8 | -3.31 |
| Risk | B*38:02_M | F | 1 | -3.10 | 9 | -4.39 | F | 1 | -3.22 | 7 | -4.39 | 2 | -3.88 |
| Possible risk | B*38:01_M | F | 0 | N/A | 10 | -3.66 | O | 6 | -3.62 | 0 | N/A | 4 | -3.89 |
| Control | B*15:01_S | B | 8 | -3.60 | 2 | -3.58 | O | 2 | -3.55 | 0 | N/A | 8 | -4.14 |
| Control | B*40:06_M | F | 0 | N/A | 10 | -3.65 | F | 0 | N/A | 10 | -3.74 | 0 | N/A |
| Control | B*46:01_M | B | 10 | -3.55 | 0 | N/A | B | 7 | -3.43 | 3 | -3.32 | 0 | N/A |
| Control | B*51:01_S | B | 5 | -3.58 | 5 | -3.54 | F | 2 | -3.54 | 8 | -3.55 | 0 | N/A |
| Control | B*54:01_M | F | 1 | -3.29 | 9 | -3.47 | F | 1 | -3.48 | 9 | -3.45 | 0 | N/A |

**Fig 3:**
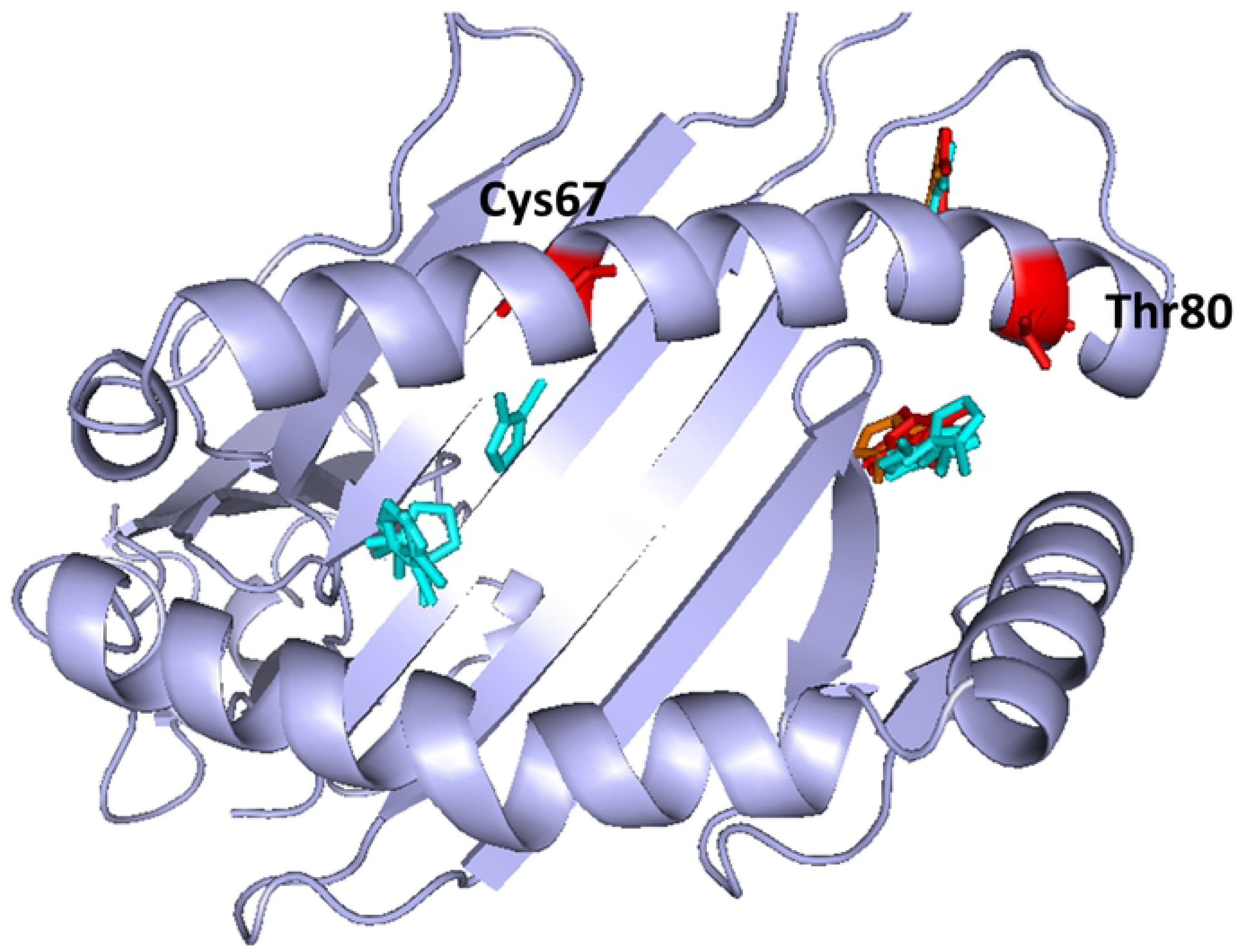
Molecular docking poses for methimazole. Top scoring docking poses of methimazole for B*38:02_M and B*27:05_S risk alleles (red), B*38:01_M non-associated allele (orange) and the control alleles (blue) B*15:01_S, B*40:06_M, B*46:01_M, B*51:01_S and B*54:01_M, using peptide binding groove search space and top3 pockets search space for AutoDockFR.

It has previously been shown that the process and parameterisation of homology modelling may have an impact on the molecular docking results, compared to docking within a crystal structure (43). It can be seen here that the control alleles favouring the F-pocket are generally the modelled structures, with the B*15:01 and B*51:01 known crystal structures showing favouring of the B-pocket. Similarly, B*38:02 and B*38:01 are both modelled structures. Due to the very high sequence similarity between these alleles, it may be that these alleles show similar binding due to the similarity of the modelled structures. Two alleles that differ little will often show similar models, although larger differences would likely have an effect. These models were shown to be structurally similar with an RMSD of 0.460Å (221 to 221 atoms).

### Propylthiouracil

Both methimazole and propylthiouracil have been associated with drug induced agranulocytosis with B*27:05 and B*38:02. Propylthiouracil was therefore also docked with the risk, possible risk and selected control alleles using AutoDockFR (39). The docking results were then compared between alleles and with the methimazole results to identify difference in favourable binding positions between the drug-allele combinations. Table 3 summarises the poses seen for propylthiouracil, showing the pocket of the lowest scoring pose, the number of poses in each pocket and the median scores for the poses in those pockets for each search space. From this and Fig 4, showing the predicted binding positions for the top scoring pose for each allele, similar patterns to those seen for methimazole can be seen. The B*27:05_S and B*38:02_M risk alleles favour the F-pocket searching the peptide binding groove and favour other pockets outside of the peptide binding groove when extending the search, with poses shown to lie close to Thr80 and B*38:02_M showing the lowest scoring pose within the F-pocket. The B*38:01_M possible risk shows favouring of the F-pocket searching both the peptide binding groove and extending to cover the top three largest pockets. Looking at the control alleles, B*46:01_M favours the B-pocket with both search spaces, B*15:01_S favours the B-pocket with scores but not with number of poses when searching the peptide binding groove and favours pockets outside of the groove when extending the search space, including a pocket close to the Thr80 position. B*40:06_M, B*51:01_S and B*54:01_M all favour the F-pocket with both search spaces. Comparing the LigPlot poses for the best scoring propylthiouracil poses (S12 Fig), searching the peptide binding groove, it can be seen that B*27:05_S, B*38:02_M, B*38:01_M, B*40:06_M, B*51:01_S and B*54:01_M all show hydrophobic interactions with propylthiouracil at the position 80 residue. B*27:05 and B*38:01 show interactions of this position with the thiocarbonyl group, B*38:02 with the carbon tail, B*54:01 with the oxygen atom and both B*40:06 and B*51:01 with one of the nitrogen atoms, with the interaction seen in B*40:06 being a hydrogen bond rather than the usual hydrophobic interaction seen.

**Table 3:** Summary of molecular docking positions for propylthiouracil. Position of the lowest scoring propylthiouracil pose along with the number of poses in each pocket and the median of the pose scores in each pocket for each of the alleles using the search space covering the peptide binding groove and the top 3 pockets identified on the protein (Extended search space). Scores given as kcal/mol. ‘O’ refers to pockets other than the B and F, i.e. outside of the binding groove.

| Status | Allele | Peptide Binding Groove |  |  |  |  | Extended Search Space |  |  |  |  |  |  |
| --- | --- | --- | --- | --- | --- | --- | --- | --- | --- | --- | --- | --- | --- |
|  |  | Lowest | B | Median | F | Median | Lowest | B | Median | F | Median | O | Median |
| Risk | B*27:05_S | F | 3 | -5.84 | 7 | -6.23 | O | 1 | -5.48 | 0 | N/A | 9 | -5.69 |
| Risk | B*38:02_M | F | 4 | -5.66 | 6 | -6.60 | F | 0 | N/A | 3 | -6.58 | 7 | -5.62 |
| Possible risk | B*38:01_M | F | 0 | N/A | 10 | -6.48 | F | 0 | N/A | 10 | -6.45 | 0 | N/A |
| Control | B*15:01_S | B | 2 | -6.34 | 8 | -6.30 | O | 0 | N/A | 0 | N/A | 10 | -6.65 |
| Control | B*40:06_M | F | 0 | N/A | 10 | -5.74 | F | 0 | N/A | 10 | -5.80 | 0 | N/A |
| Control | B*46:01_M | B | 10 | -6.43 | 0 | N/A | B | 10 | -6.41 | 0 | N/A | 0 | N/A |
| Control | B*51:01_S | F | 0 | N/A | 10 | -6.45 | F | 3 | -6.00 | 7 | -6.44 | 0 | N/A |
| Control | B*54:01_M | F | 0 | N/A | 10 | -6.42 | F | 0 | N/A | 10 | -6.41 | 0 | N/A |

**Fig 4:**
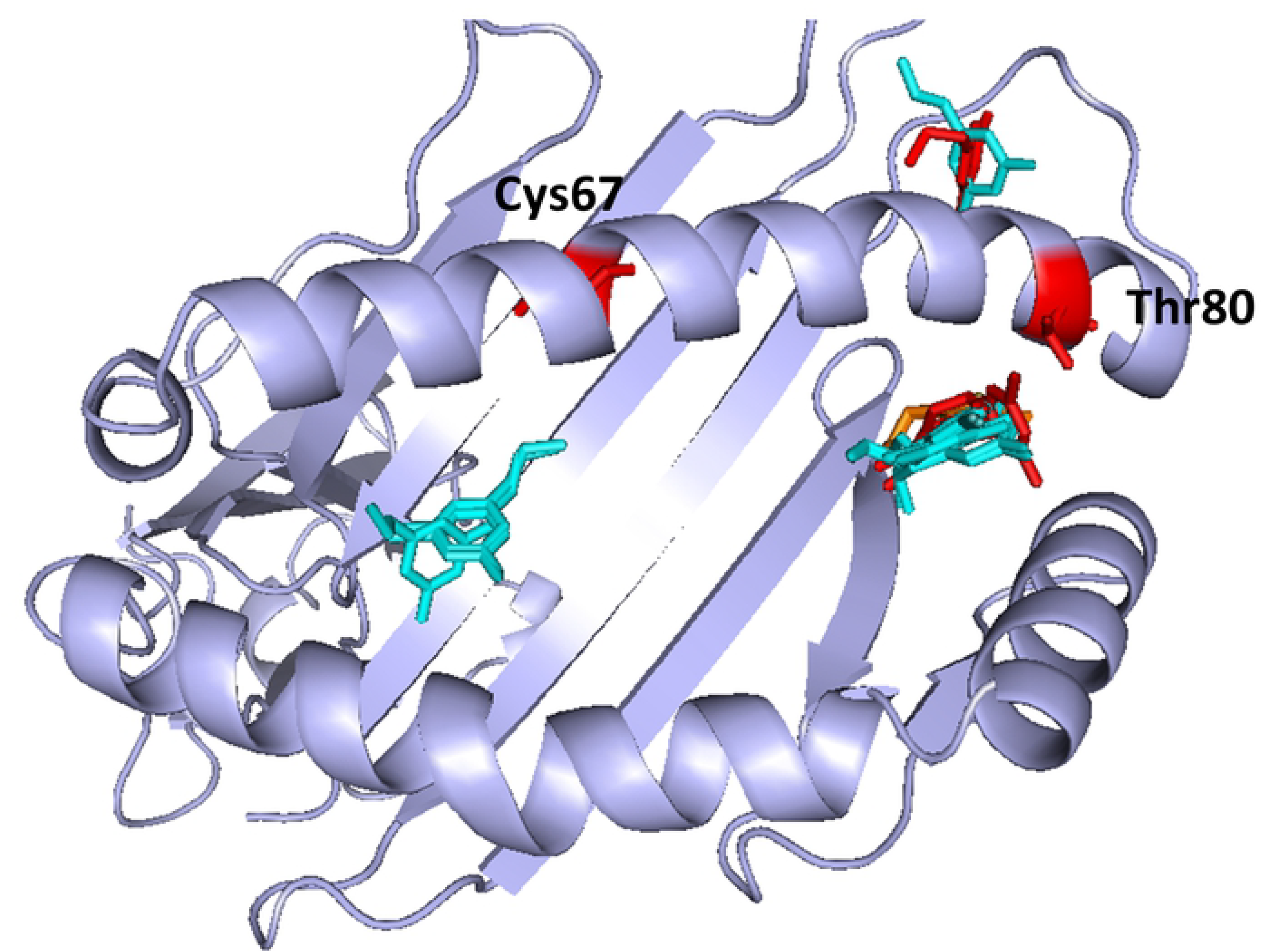
Molecular docking poses for propylthiouracil. Top scoring docking poses of propylthiouracil for B*38:02_M and B*27:05_S risk alleles (red), B*38:01_M non-associated allele (orange) and the control alleles (blue) B*15:01_S, B*40:06_M, B*46:01_M, B*51:01_S and B*54:01_M, using peptide binding groove search space and top3 pockets search space for AutoDockFR.

Fig 5 shows the docking scores for all drug-allele combinations, searching the peptide binding groove, when expanding the analysis to 100 runs. From this we can see consistent favouring of the F pocket for the risk alleles. Although the docking is not able to fully distinguish between the risk and controls, with the controls showing mixed favouring of the B and F pockets, it is evident that docking against the risk alleles strongly favours binding in the F pocket for each of the drugs.

**Fig 5:**
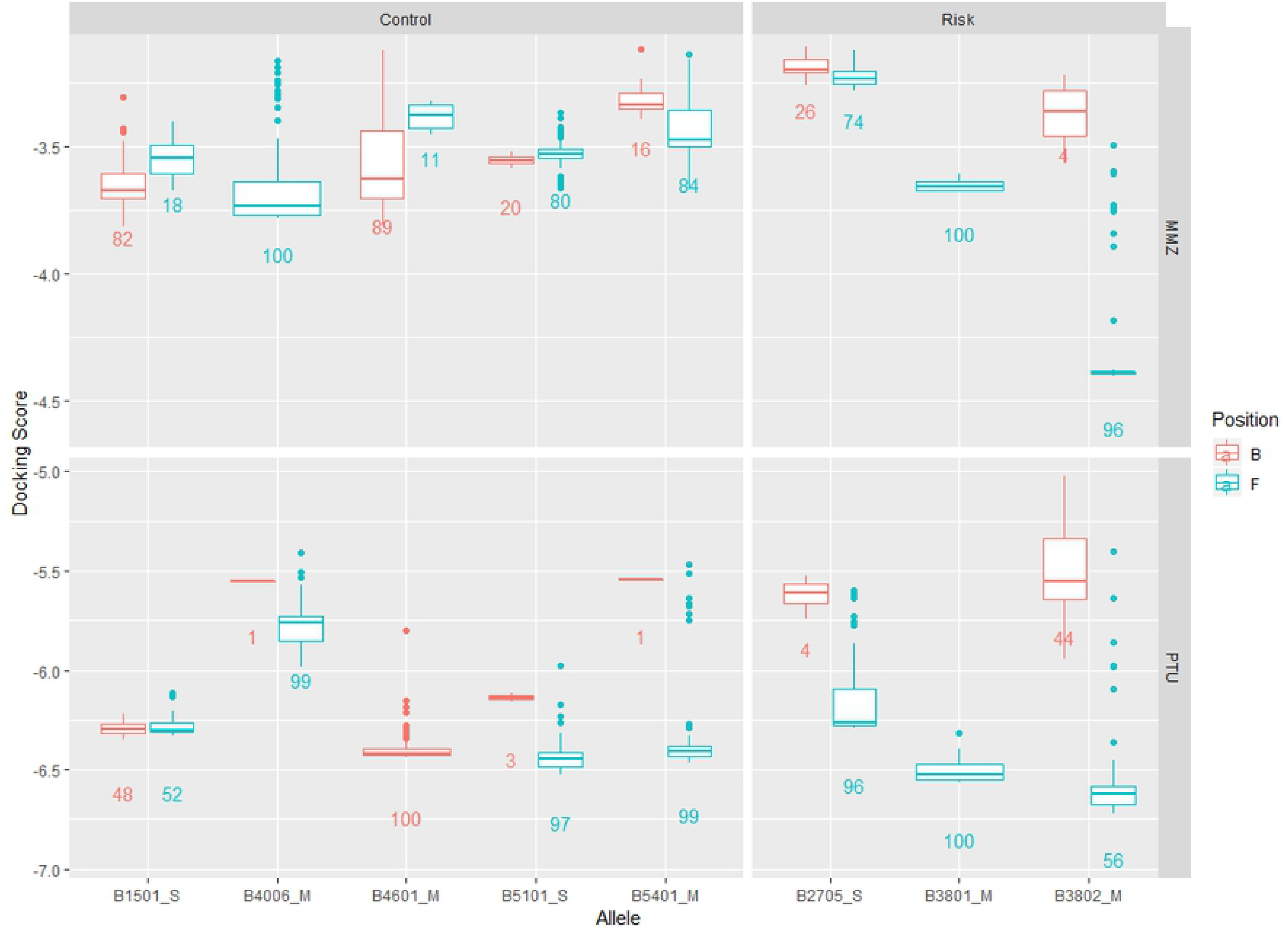
Boxplots for methimazole and propylthiouracil. Boxplots showing docking scores for 100 poses searching the peptide binding groove, using both methimazole (MMZ) and propylthiouracil (PTU) for each of the alleles.

Similar molecules to the investigated methimazole and propylthiouracil were identified: MZY, TUL, DMI and EV0. The similar molecules were docked to each of the risk and control alleles in order to deduce if the size and structures of the ligands and pockets could be having an impact on the molecular docking results (S5 Fig). From these investigations, it was found that the similar molecules that have been used as experimental anti-thyroid drugs and included the thiocarbonyl group (MZY and TUL), showed similar binding patterns to the associated anti-thyroid drugs with the risk alleles favouring the F-pocket. The possible risk B*38:01_M allele can be seen to favour the B-pocket by scores but the F by number of poses for MZY and favour the F for TUL (S2 Table). Control alleles B*15:01_S and B*46:01_M both favour the B via lower scores and more poses for both drugs. With control B*51:01_S also favouring the B for TUL considering scores but not poses. For the other molecules without the sulfhydryl group (DMI and EV0), a difference in trends is seen with the B*27:05_S favouring the B-pocket for both drugs and B*38:02_M showing the lowest scoring pose for DMI favouring the B-pocket but still favouring the F based on poses and for EV0. B*38:01_M possible risk still shows favouring of the F-pocket for these drugs. The controls all show favouring of the B through scores and number of poses for both drugs, except B*54:01_M, which shows favouring of the F-pocket for EV0. Docking poses are shown in S13 Fig.

Table 4 summarises the number of poses making hydrophobic interactions with position Thr80 of both risk alleles for each of the investigated drugs. It can be seen that for B*27:05 the associated drugs often make Thr80 interactions with interactions with the thiocarbonyl group being most favourable. The experimental drugs show similar interactions with Thr80 as those seen with the associated drugs (S14 Fig). For B*38:02, fewer interactions are made between the drugs and the position 80 Thr residue than seen for B*27:05 (S15 Fig). In both risk alleles, the most favourable poses commonly form interactions between the thiocarbonyl group of methimazole, propylthiouracil, MZY and TUL, and positions 77, 80, 81, 84 and 123. Interactions are also seen with positions 95, 116, 124, 143 and 147. All these residues seen making interactions surround the F-pocket. Positions 77, 80 and 123 were identified as of interest by the Chen *et al.* study (18). This adds strength to the hypothesis that the Thr80 position could be involved in the mechanism here, especially for B*27:05. The structure of the ligands, mainly the ‘S’ group found in the associated and experimental anti-thyroid drugs, could therefore be potentially important for the predicted binding poses seen here.

**Table 4:** Summary of Thr80 interactions for investigated ligands. Summary counts showing the number of poses seen making interactions with the Thr80 residue of each of the associated risk alleles for the associated drugs (MMZ and PTU), the experimental anti-thyroid drugs containing the thiocarbonyl group (MZY and TUL) and the other investigated ligands. Where ‘No Thr 80’ relates to poses showing no interactions between the Thr80 residue and the drug, ‘Thr80-S’ shows interactions between the thiocarbonyl group and Thr80 and ‘Other Thr80’ indicates interactions made between Thr80 and the drug but not with the thiocarbonyl group. The ‘Most favourable pose’ column shows the interaction seen for the pose which showed the lowest docking score for each of the drug-allele combinations.

| Drug | B*27:05 |  |  |  | B*38:02 |  |  |  |
| --- | --- | --- | --- | --- | --- | --- | --- | --- |
|  | No Thr80 | Thr80-S | Other Thr80 | Most favourable pose | No Thr80 | Thr80-S | Other Thr80 | Most favourable pose |
| MMZ | 5 | 4 | 1 | Thr80-S | 10 | 0 | 0 | No Thr80 |
| PTU | 3 | 5 | 2 | Thr80-S | 5 | 2 | 3 | Other Thr80 |
| MZY | 1 | 2 | 7 | Other Thr80 | 9 | 0 | 1 | No Thr80 |
| TUL | 5 | 2 | 3 | Thr80-S | 10 | 0 | 0 | No Thr80 |
| DMI | 9 | 0 | 1 | No Thr80 | 10 | 0 | 0 | No Thr80 |
| EV0 | 10 | 0 | 0 | No Thr80 | 5 | 0 | 5 | Other Thr80 |

## Discussion

The purpose of this study was to investigate the associations seen between HLA and anti-thyroid alleles, focusing on the commonalities seen between the HLA-B associated alleles, to identify a potential shared mechanism. This was done through comparing the peptide binding regions, including the whole peptide binding groove and specific residue changes alongside the predicted binding positions of the drugs with each of the risk and control alleles. It was found that the risk alleles favour different peptides and so we can conclude that the gross structures of their peptide binding grooves are rather dissimilar. However, when the multiple sequence alignments were used to focus on specific residue changes, it could be seen that two residues were found to be unique to the risk alleles. These two residues, Cys67 and Thr80, were therefore considered to be potentially important for the mechanism of action for the adverse drug reactions seen, confirming the results of a previous study by Chen *et al*. where the Cys67 and Thr80 were identified, amongst others, as potentially important for the binding of the associated drugs with the risk alleles (18). From the results of the molecular docking, the risk alleles were shown to favour the F-pocket for both drugs. This pocket lies alongside the Thr80 residue which was identified as potentially important for the mechanism of action. It was seen that the Thr80 interacts hydrophobically with the thiocarbonyl group of both associated drugs in B*27:05, with similar interactions also being seen, but to a lesser extent, with B*38:02. The residue at position 80 of the control alleles was seen to only make interactions with the methimazole for B*54:01, although these were different interactions from those seen for the risk alleles. For propylthiouracil, this position made interactions with the molecule for three of the five control alleles, although these were again seen as different interactions to the risk, with the B*38:01 possible risk showing similar interactions to the B*27:05 risk allele. It is therefore reasonable to conclude that Thr80 could be involved in the mechanism of interaction, if only indirectly by influencing the conformations of the surrounding residues located around the F-pocket.

Although the docking results for the risk alleles showed consistent results, with the risk alleles always favouring the F-pocket, the predicted poses for the control alleles showed some variation with some drug-allele combinations favouring the B-pocket and some favouring the F-pocket. Docking has previously been shown to be imperfect when considering these complex HLA cases (43), it is therefore understandable that the docking was unable to distinguish completely between risk and control for this case. Docking results are commonly reported as the most favourable pose, here we show a comparison of multiple runs as well as comparisons with selected control alleles The risk alleles are shown to continuously favour the F-pocket over many runs, providing evidence that this is more likely position of binding. The variation seen between controls could be due to a number of factors including the homology modelling of the protein structures. As discussed in a previous study (43), homology modelling can impact the docking performance and inaccurate models could result in inaccurate docking results. Here, the similar alleles B*38:02 (risk) and B*38:01 (possible risk), with one mutated residue at position 80, were docked with both of the associated drugs. These alleles showed similar binding patterns with both alleles generally favouring the F-pocket. Since we have concluded here that the Thr80 could potentially be involved in the mechanism of action, it would be expected that the B*38:01 would show different docking results to the risk alleles as this allele does not possess a Threonine residue at position 80. However, both the B*38:02 and B*38:01 structures were obtained through homology modelling. This could impact the results of the molecular docking as although the mutation has been modelled, it may not have been accurately represented here and the similar structures could produce similar models.

The results seen here confirm and build on those seen in the Chen *et al.* study (18), with Thr80 being identified as important for the mechanism of the adverse drug reaction. The Chen *et al*. study conducted docking of B*38:02, B*38:01 and DRB1*08:03 with methimazole and propylthiouracil. Models were created for B*38:02 and B*38:01 using the same five templates (A*24:02, C*08:01 and A*02:01 along with mouse MHC and human HLA-E) and the model for DRB1*08:03 created using one template (DRB1*01:01). The templates used for the B alleles show similarities of 83-86% with B*38:02 and the DRB1*01:01 template shows 92% identity with DRB1*08:03. In this study, the B*38:02 model was created using templates with 95-98% identity with differing templates with similar identity being used for the other modelled structures created. Model selection is a very important aspect of molecular docking and can greatly impact the docking results seen (43). This study was able to recreate similar docking results seen previously for B*38:02 and B*38:01, using our own modelled structures, whilst also incorporating docking results for the B*27:05 risk allele and comparisons with selected control alleles. Here, we also went further to investigate the structural differences between the associated risk alleles and selected controls, through sequence alignments and comparison of binding motifs.

In order to further our understandings of the mechanisms involved here and to test the hypothesis that the Thr80 is important for binding, it would be interesting to investigate B*38:01 further as this is a good potential biological control due to its similarity with the associated B*38:02. For this, further association studies to confirm the association status of B*38:01 would be needed. Structural analysis through crystal structures would be needed to confirm the binding predictions of the drug allele combinations investigated and therefore the involvement of Thr80 in the predisposition to this adverse drug reaction.

## Acknowledgments

The research was part-funded by the Medical Research Council grant for the Centre for Drug Safety Science, University of Liverpool (Grant Number: MR/L006758/1). The funders had no role in study design, data collection and analysis, decision to publish, or preparation of the manuscript.

## Supporting Information

***S1 Table: Structures obtained for risk and control alleles***

***S2 Table: Summary of molecular docking positions for investigated compounds.*** *Position of the lowest scoring poses for each of the drug-allele combinations, along with the number of poses in each pocket and the median of the pose scores in each pocket for each of the alleles using a search space covering the peptide binding groove. Scores given as kcal/mol.*

***S1 Fig: Bar chart of allele frequencies for investigated populations.*** *Healthy population frequencies obtained from AFND (24) along with Case and Control frequencies for alleles from the He et al. study (17) (Han Northern China) with control frequency over 3% and alleles from the Hallberg et al. study (20) with AFND frequency greater than 3% in at least one population (Sweden, France, Spain, or Germany). Panels show alleles separated not only by population but also separating those alleles that were selected as suitable controls.*

***S2 Fig: Clustering analysis and docking predicted docking poses of 100 runs.*** *F pocket predictions shown in purple, B pocket predictions shown in yellow. Top scoring docking poses of methimazole and propylthiouracil for B*38:02_M and B*27:05_S risk alleles (red), B*38:01_M non-associated allele (orange) and the control alleles (blue) B*15:01_S, B*40:06_M, B*46:01_M, B*51:01_S and B*54:01_M, using peptide binding groove search space for AutoDockFR.*

***S3 Fig: Alignment of positions potentially involved in binding for the B and F pockets.*** *Unique matches identified at these positions are shown highlighted in red.*

***S4 Fig: Organisation of subsites along the HLA peptide binding groove.*** *The six subsites along the peptide biding groove (A-F) are shown highlighted (44, 45). Image created using PyMOL (46).*

***S5 Fig: Structure of investigated drugs.*** *Structure of associated drugs methimazole and propylthiouracil along with similar ligands, some of which have been used as experimental anti-thyroid drugs.*

***S6 Fig: Allele frequency distribution for B*38:02.*** *Allele frequency distribution map obtained from AFND (24). The size of the circles represent the sample size of each population with the colour representing the allele frequency, as shown by the key, with low frequency alleles shown in blue, mid in green and high in orange/red.*

***S7 Fig: Allele frequency distribution for B*38:01.*** *Allele frequency distribution map obtained from AFND (24). The size of the circles represent the sample size of each population with the colour representing the allele frequency, as shown by the key, with low frequency alleles shown in blue, mid in green and high in orange/red.*

***S8 Fig: Allele frequency distribution for B*27:05.*** *Allele frequency distribution map obtained from AFND (24). The size of the circles represent the sample size of each population with the colour representing the allele frequency, as shown by the key, with low frequency alleles shown in blue, mid in green and high in orange/red.*

***S9 Fig: MHC motif viewer output for risk and control alleles.*** *MHC Motif Viewer outputs for B*27:05 and B*38:02 risk alleles and B*38:01 possible risk alleles, showing predicted peptide binding motifs for each allele. Amino acids are coloured according to their physicochemical properties; acidic (D, E) coloured red, basic (H, K, R) coloured blue, hydrophobic (A, C, F, I, L, M, P, V, W) coloured black and Neutral (G, N, Q, S, T, Y) coloured green. The height of the column of letters is equal to the information content at that position and the height of the letter within the column is proportional to the frequency of the corresponding amino acid at that position. Where available, the reliability index is shown in the centre of the circle above the logo plot and is given as the estimated Pearson correlation coefficient for neural network predictions on the given alleles. The closest neighbour is also shown along with the distance to this neighbouring allele (25).*

***S10 Fig: Multiple sequence alignment for risk, control and common alleles.*** *(a) Multiple sequence alignment for risk alleles B*27:05 and B*38:02, possible risk allele B*38:01 and top 10 most frequent alleles for Caucasian, Asian populations selected from AFND and NMDP populations (repeats removed). (b) Focusing on positions 65-83. (c) Multiple sequence alignment for risk alleles B*27:05 and B*38:02, along with possible risk allele B*38:01 and AFND top 20 most common alleles and NMDP top 10 most common for Caucasian, Asian and all populations (repeats removed – 30 unique sequences including risk alleles). (d) Focusing on positions 65-83. Positions 67 and 80 highlighted in red.*

***S11 Fig: LigPlot figures for methimazole.*** *LigPlot figures show the interactions between methimazole and the residues on each allele for each of the risk, suspected risk and control alleles, for the top scoring pose searching the peptide binding groove. Circles show comparisons between alleles, amino-acids at positions found interacting for B*27:05 highlighted with red circle. Hydrogen bonds are shown by green dotted lines.*

***S12 Fig: LigPlot figures for propylthiouracil.*** *LigPlot figures show the interactions between propylthiouracil and the residues on each allele for each of the risk, possible risk and control alleles, for the top scoring pose searching the peptide binding groove. Circles show comparisons between alleles, amino-acids at positions found interacting for B*27:05 highlighted with red circle. Hydrogen bonds are shown by green dotted lines.*

***S13 Fig: Predicted binding poses for investigated ligands.*** *Top scoring binding poses for B*27:05_S searching the peptide binding groove for each of the investigated drugs. Red poses show associated allele poses, orange shows B*38:01_M non-associated allele poses and blue the control allele poses.*

***S14 Fig: LigPlot figures for B*27:05.*** *LigPlot figures show the interactions between B*27:05 and each of the investigated ligands for the top scoring pose searching the peptide binding groove. Circles show comparisons between alleles, amino-acids at positions found interacting for MMZ highlighted with red circle. Hydrogen bonds shown by green dotted lines. Hydrophobic bonds shown in red.*

***S15 Fig: LigPlot figures for B*38:02.*** *LigPlot figures show the interactions between B*38:02 and each of the investigated ligands for the top scoring pose searching the peptide binding groove. Circles show comparisons between alleles, amino-acids at positions found interacting for MMZ highlighted with red circle. Hydrogen bonds shown by green dotted lines. Hydrophobic bonds shown in red.*

